# Genomic characteristics of recently recognized *Vibrio cholerae* El Tor lineages associated with cholera in Bangladesh, 1991-2017

**DOI:** 10.1101/2021.12.16.473094

**Authors:** Md Mamun Monir, Talal Hossain, Masatomo Morita, Makoto Ohnishi, Fatema-Tuz Johura, Marzia Sultana, Shirajum Monira, Tahmeed Ahmed, Nicholas Thomson, Haruo Watanabe, Anwar Huq, Rita R. Colwell, Kimberley Seed, Munirul Alam

**Affiliations:** icddr, b, Formerly International Centre for Diarrhoeal Disease Research, Bangladesh, Dhaka, Bangladesh; National Institutes of Infectious Diseases (NIID), Tokyo, Japan; Sanger Institute, Cambridge, UK; Maryland Pathogen Research Institute, University of Maryland, USA; Johns Hopkins Bloomberg School of Public Health, Baltimore, Maryland, USA; University of California, Berkeley, USA

## Abstract

Comparative genomic analysis of *Vibrio cholerae* El Tor associated with endemic cholera in Asia revealed two distinct lineages, one dominant in Bangladesh and the other in India. An in depth whole genome study of *V. cholerae* El Tor clinical strains isolated during endemic cholera in Bangladesh (1991 – 2017) included reference genome sequence data obtained online. Core genome phylogeny established using single nucleotide polymorphisms (SNPs) showed *V. cholerae* El Tor strains comprised two lineages, BD-1 and BD-2, which, according to Bayesian phylodynamic analysis, originated from paraphyletic group BD-0 around 1981. BD-1 and BD-2 lineages overlapped temporally but were negatively associated as causative agents of cholera 2004-2017. Genome wide association study (GWAS) revealed 140 SNPs and 31 indels, resulting in gene alleles unique to BD-1 and BD-2. Regression analysis of root to tip distance and year of isolation indicated early BD-0 strains at the base, whereas BD-1 and BD-2 subsequently emerged and progressed by accumulating SNPs. Pangenome analysis provided evidence of gene acquisition by both BD-1 and BD-2, of which six crucial proteins of known function were predominant in BD-2. BD-1 and BD-2 diverged and have distinctively different genomic traits, namely heterogeneity in VSP-2, VPI-1, mobile elements, toxin encoding elements, and total gene abundance. In addition, the observed phage-inducible chromosomal island-like element (PLE1), and SXT ICE elements (ICE^TET^) in BD-2 presumably provided a fitness advantage for the lineage to outcompete BD-1 as the etiological agent of the endemic cholera in Bangladesh, with implications for global cholera epidemiology.

**Importance:** Cholera is a global disease with specific reference to the Bay of Bengal Ganges Delta where *Vibrio cholerae* O1 El Tor, causative agent of the disease showed two circulating lineages, one dominant in Bangladesh and the other in India. Results of in-depth genomic study of *V. cholerae* associated with endemic cholera during the past 27 years (1991 – 2017) indicate emergence and succession of the two lineages, BD-1 and BD-2, arising from a common ancestral paraphylatic group, BD-0, comprising the early strains and short-term evolution of the bacterium in Bangladesh. Among the two *V. cholerae* lineages, BD-2 supersedes BD-1 and is predominant in the most recent endemic cholera in Bangladesh. The BD-2 lineage contained significantly more SNPs and indels, and showed richness in gene abundance, including antimicrobial resistance genes, gene cassettes, and PLE to fight against bacteriophage infection, acquired over time. These findings have important epidemic implications at a global scale.

## Introduction

Cholera is a life threatening infectious diarrheal disease caused by *Vibrio cholerae* serogroups O1 and O139 of the Gram-negative gammaproteobacteria (1, 2). The global incidence of cholera is estimated to be 2.9 million cases annually with almost 95,000 deaths (3). In 2017, 34 countries reported a total of 1,227,391 cases and 5,654 deaths (4). Seven cholera pandemics have been recognized since 1817. However, limited information is available regarding the etiological agent for the first five pandemics and no isolates of the causative agent are extant. The sixth pandemic, and possibly those earlier were caused by *V. cholerae* O1 classical biotype, while the ongoing seventh pandemic is caused by *V. cholerae* El Tor biotype and began with displacement of *V. cholerae* classical biotype in Asia in 1961 (5). *V. cholerae* El Tor was isolated in Africa in the 1970s and Latin America in 1991 where for more than a century there had been no cholera outbreaks (6). In 1992, a *V. cholerae* non-O1 strain designated *V. cholerae* O139 Bengal initiated outbreaks of cholera in coastal areas of India and Bangladesh, and subsequently was isolated from patients in several countries of Asia (2). *V. cholerae* El Tor continues to be the major etiological agent of cholera worldwide.

The severe dehydrating diarrhea characteristic of cholera is associated with several factors, including a toxin and several virulence genes involved in colonization and toxicity and their coordinated expression (1). Cholera toxin (CT) is the virulence factor responsible for secretory diarrhea of cholera and is encoded in the genome of a lysogenic CTX phage. *V. cholerae* El Tor responsible for the current cholera pandemic harbors the CTX phage classical biotype variant, and the ctxB^cla^: ctxB genotype 1 (*ctxB*1) or *ctxB*7 (7). *V. cholerae* responsible for the current cholera pandemic has become more virulent by undergoing several shifts in CTX genotype and acquiring virulence-related gene islands (8). Integrative conjugative elements (ICEs) and lysogenic phages are genetic elements that play an important role in the acquisition of virulence, antimicrobial resistance, and heavy metal resistance, which important components of the pathogenicity of *V. cholerae* (9, 10). Functions of these elements are important for the pathogen to exert evolutionary advantage and variants can be used as markers of clonal expansion (1). Acquisition of mobile genetic elements (MGEs) through horizontal gene transfer (HGT) and propitious chromosomal mutations are significant landmarks for an evolving bacterium (11).

Whole-genome sequencing of *V. cholerae* El Tor strains associated with the seventh cholera pandemic revealed three waves, suggesting independent but overlapping paths for the pathogen to spread globally from the Bay of Bengal estuary where cholera has been endemic at least since 1961 but likely for centuries (5). Intercontinental transmission of *V. cholerae* has been proposed for the 2010 outbreak in Haiti (12). Bangladesh borders on the Bay of Bengal and is considered to be a hotspot of Asiatic cholera where -ca. 100,000 cases and 4,500 deaths are reported each year (13). *V. cholerae* O1 responsible for endemic cholera in Bangladesh and India has been found to have undergone genetic changes over time including acquisition of classical biotype attributes in an El Tor background, thereby becoming more successful as a pathogen (14, 15). A recent whole-genome analysis of *Vibrio cholerae* El Tor strains isolated between 2009 and 2016 indicated two distinct lineages exist in Bengal (16). The objective of the study reported here was to investigate *V. cholerae* endemic cholera strains isolated during 1991 to 2017 to understand more completely about emergence and progression of the two lineages in Bangladesh. Virulence and related genomic islands, including toxin and antimicrobial resistance genes differing significantly among the *V. cholerae* El Tor lineages, were also investigated for potential relevance to emergence of the lineages.

## Results

### Phylogenetic analysis

A total of 119 strains were included in the study and their genomes were sequenced using the Illumina platform (MiSeq or HiSeq 2500 sequencer). In addition, 56 strains from our previous study (16) and 17 genomes from the European Nucleotide Archive (17) were used, which are representative of isolates from Bangladesh between 1991 and 2017 (see Table S1 Supplemental Material). Paired-end reads of the 192 genomes were mapped to *V. cholerae* El Tor N16961 reference strain, a seventh-pandemic *V. cholerae* O1 El Tor (7PET) strain isolated in Bangladesh in 1975 (18). A total of 1,298 single nucleotide polymorphisms (SNPs) and 413 indels (insertions or deletions) were obtained and, after filtering indels, low call rate, and high-density SNPs, a total of 893 high-quality SNPs were retained for further study. A phylogenetic analysis was conducted to construct a tree based on the 893 high-quality SNPs to evaluate the genetic diversity of the *Vibrio cholerae* O1 El Tor isolates from Bangladesh. A nested hierarchical structure in the phylogenetic tree was observed, with all but four of the strains isolated between 1999 and 2017 clustering into two major clades, BD-1 (n=76) and BD-2 (n=105), shown in green and red, respectively. The remaining strains formed paraphyletic group BD-0 (n=11) (Fig. 1A, in blue). Except for three strains isolated in 2012 that formed a sub-clade, BD-0 consisted mostly of strains isolated earlier between 1991 and 2000. Dates of isolation of common ancestors of the lineages were inferred using Bayesian Markov chain Monte Carlo framework Bayesian Evolutionary Analysis Sampling Trees (BEAST) (19) (see Fig. S1 supplementary material), and a maximum clade credibility (MCC) tree was inferred from the posterior distribution of the best fitting model using program TreeAnnotator tool of the BEAST software package. It was estimated from the MCC tree that the most recent common ancestor (MRCA) of lineage BD-1 was isolated in 1987 (95% HPD: 1983-1991), and lineage BD-2 in 1997 (95% HPD: 1994-2000), where HPD stands for height posterior density. Strains of BD-1 and BD-2 shared genome sequences of strains isolated since 1981 (95% HPD: 1976-1986). The number of SNPs in strains of the two clades is relative to reference *V. cholerae* N16961, which showed strains of BD-0 differed by 107 - 137 SNPs, BD-1 by 123 - 189 SNPs, and BD-2 by 146 - 186 SNPs. An unrooted tree showed SNP diversity among BD-0, BD-1, and BD-2 clades with SNP diversity of BD-2 highest (Fig. 1B). Comparison of isolates in the clades and year of isolation revealed clonal aggregation within the dominant clade and strong temporal signature. Strains of BD-1 and BD-2 were found to be temporally spread but simultaneously isolated during the periods of 2004 - 2011, 2012, 2014 - 2016 (Fig. 1C, Table S2 supplemental material). Strains of BD-1 were mainly isolated during 2004-2011 (66.3%, n=65) while strains of BD-2 were isolated during those years in fewer numbers (33.7%, n=33) except 2009 when BD-2 strains were dominant (93.33%, n=14) (see Table S1 in supplementary material). The following years, from 2012 to 2017, showed BD-2 strains to be dominant (73.5%, n=72) and BD-1 strains the minority (10.2%, n=10).

**FIG 1.**
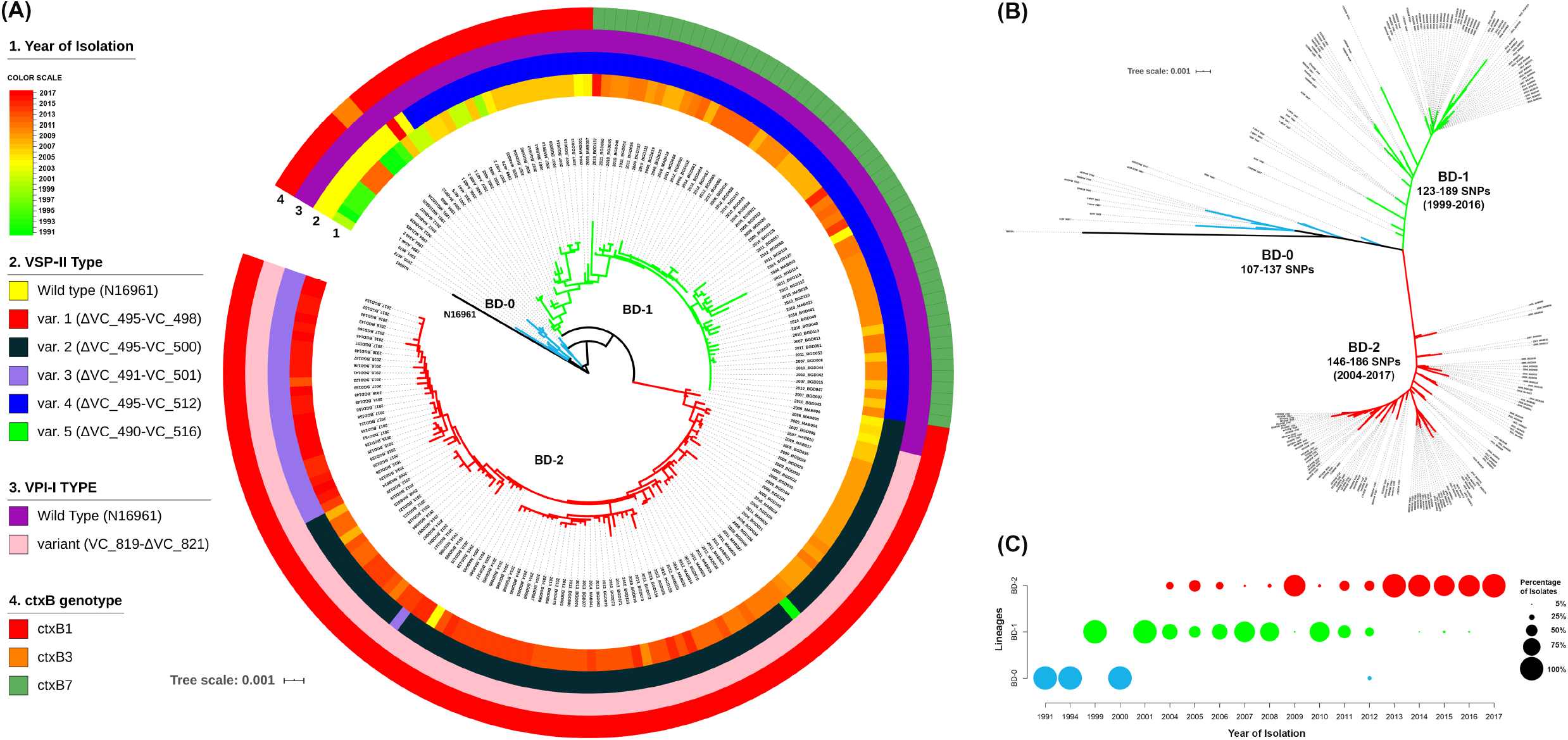
Phylogenetic analyses of strains showing respective genomic features and year of isolation. (A) Maximum likelihood phylogenetic tree generated from whole genome SNPs and number of isolated *V. cholerae* O1 El Tor strains belonging to lineages BD-0, BD-1, and BD-2 rooted from out-group reference strain *Vibrio cholerae* N16961. Rings show features of the isolates according to color scheme provided on the left. Tree branches are colored blue, green, and red defining lineages BD-0, BD-1, and BD-2, respectively; (B) Unrooted tree showing independent evolution of BD-1 and BD-2 strains with the number of core genome SNPs of strains in the lineages compared to the N16961 reference strain; and (C) Percentage of isolates per year for the three lineages. Size of the circles indicates percentage of strains belonging to lineages according to the scheme shown.

### Genetic variants associated with the clades

Associations between lineages and the genetic variants was studied using 1298 SNPs and 413 indels, identified by aligning raw reads against *V. cholerae* N16961 reference genome. Variant annotation using SnpEff (20) showed that among the 1298 SNPs, there were 337 synonymous, 613 nonsynonymous, and 348 variants on intergenic regions (Fig. 2A-C, see Table S2 in the supplemental materials). Moreover, of 413 indels, there were 238 frameshift-variants, 107 variants on intergenic regions, and 68 other types of variants (Fig 2D-F, Table S2). Most of the identified SNPs and indels were located in the protein-coding region, many of which function to change the form of a protein. By plotting distribution of SNP types and indel variants for BD-0 (n=11), BD-1 (n=76), and BD-2 (n=105), it was observed that strains of the clades accumulated SNPs and indels. Strains of BD-2 accumulated more SNPs and indels, increasing genetic distance from BD-0 and BD-1 (Fig. 1B, Fig. 2) and suggesting evolution was occurring when compared with reference *V. cholerae* O1 N16961.

**FIG 2.**
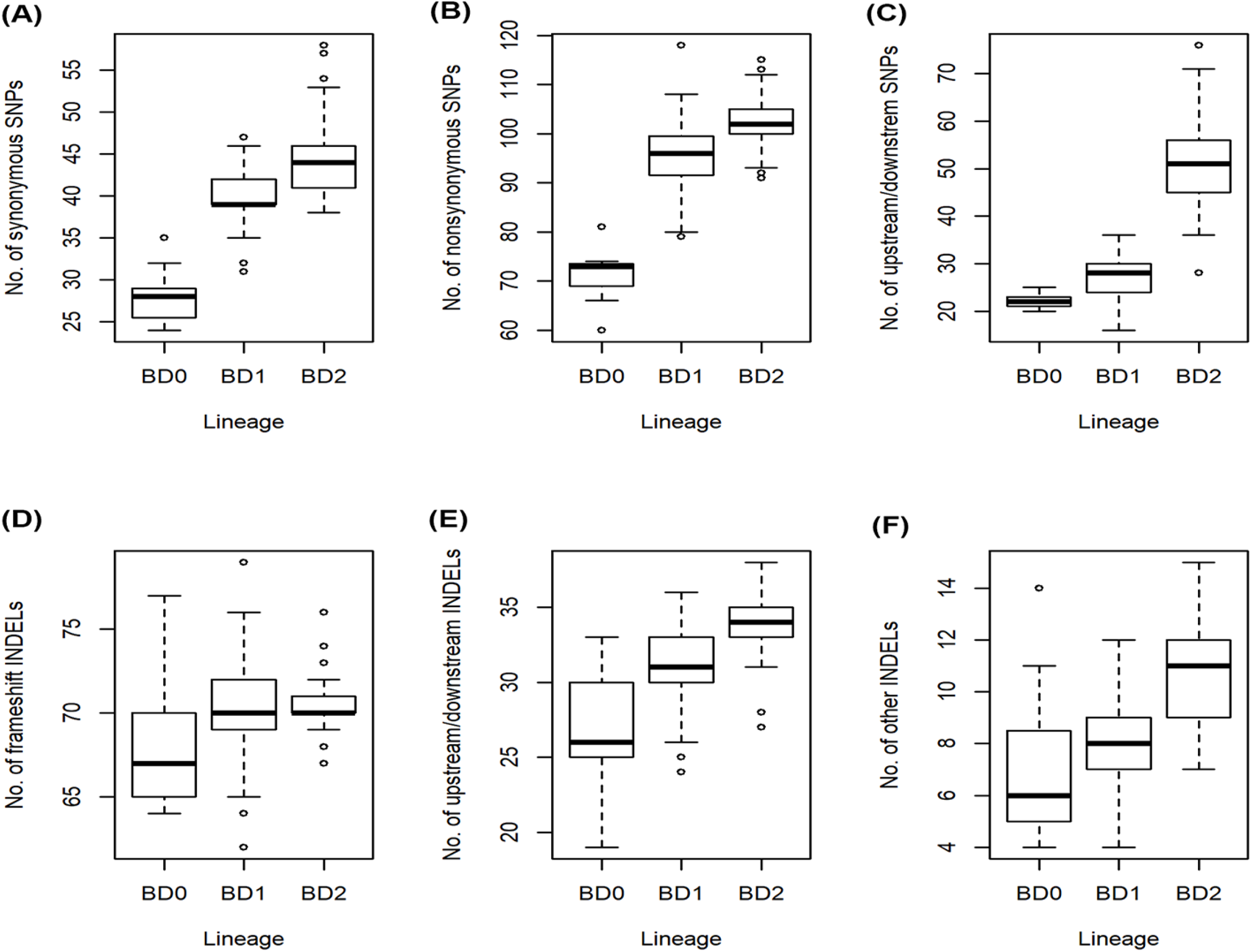
Box plots of SNPs distribution and indel type in each of three lineage groups. (A) Distribution of 337 synonymous SNP variants. This figure shows that strains of BD-2 lineage accumulated more synonymous SNP variants compared to BD-0 and BD-1 lineages. Notably, synonymous SNP variants do not change the form of protein. (B) Distribution of 613 nonsynonymous SNP variants. These nonsynonymous SNP variants include 570 missense variants, 38 stop gained variants, 2 splice-region-variants and stop-retained-variants, 2 stop-lost and splice-region-variants, 1 initiator codon variant. (C) Distribution of 348 upstream/downstream SNP variants. (D) Distribution of 238 frameshift indel variants. (E) Distribution of 107 upstream/downstream indel variants. (F) Distribution of 68 indel variants, including 13 conservative-inframe-insertions, 14 disruptive-inframe-insertions, 11 frameshift-variant and stop-gained, 10 disruptive-inframe-deletions, 10 conservative-inframe-deletions, 1 stop-gained and disruptive-inframe-deletions, 2 feature-elongations, 1 frameshift-variant and stop-lost and splice-region-variant, 1 stop-gained and disruptive-inframe-insertion, 2 frameshift-variant and splice-region-variant, 2 frameshift-variant and start-lost, 1 stop-gained and conservative-inframe-insertion.

Fisher exact test (21) was performed for association analysis between genetic variants and the clades BD-1 and BD-2. Association analysis showed that 140 SNPs and 31 indels had a genomewide significant association (*p* < 6.40×10^−9^) with BD-1 and BD-2. Among the 140 SNPs were 25 synonymous variants, 53 missense variants, 2 stop gain variants, and 60 variants on intergenic regions (Table S3 and Fig. S2 in supplementary material). It was discovered that 21 SNP missense mutations were present in genes with known functions in more than 80% of BD2 strains, resulting in mutant proteins (Table 1). However, there were only seven missense mutations were found in genes with known functions in more than 80% of BD1 strains. Genotype and frequency of 140 significantly associated SNPs, number of SNPs by year of isolation, and root to tip distance, showed significant genetic differences between BD-1 and BD-2 (Fig. 3). The number of core genome SNPs by year of isolation was analyzed to detect temporal SNP accumulation patterns of the clades. The number of core genome SNPs did increase over time for both BD-1 and BD-2 (Fig. 3B). Moreover, root-to-tip regression analysis indicated a steady increase in SNP divergence among the strains of the two clades over time (Fig. 3C). Miami plot for frequency of alternative alleles of the 140 significant SNPs showed BD-2 strains had accumulated more clade specific SNPs, notably in chromosome-2 compared to BD-1 (Fig. 3D).

**Table 1.**
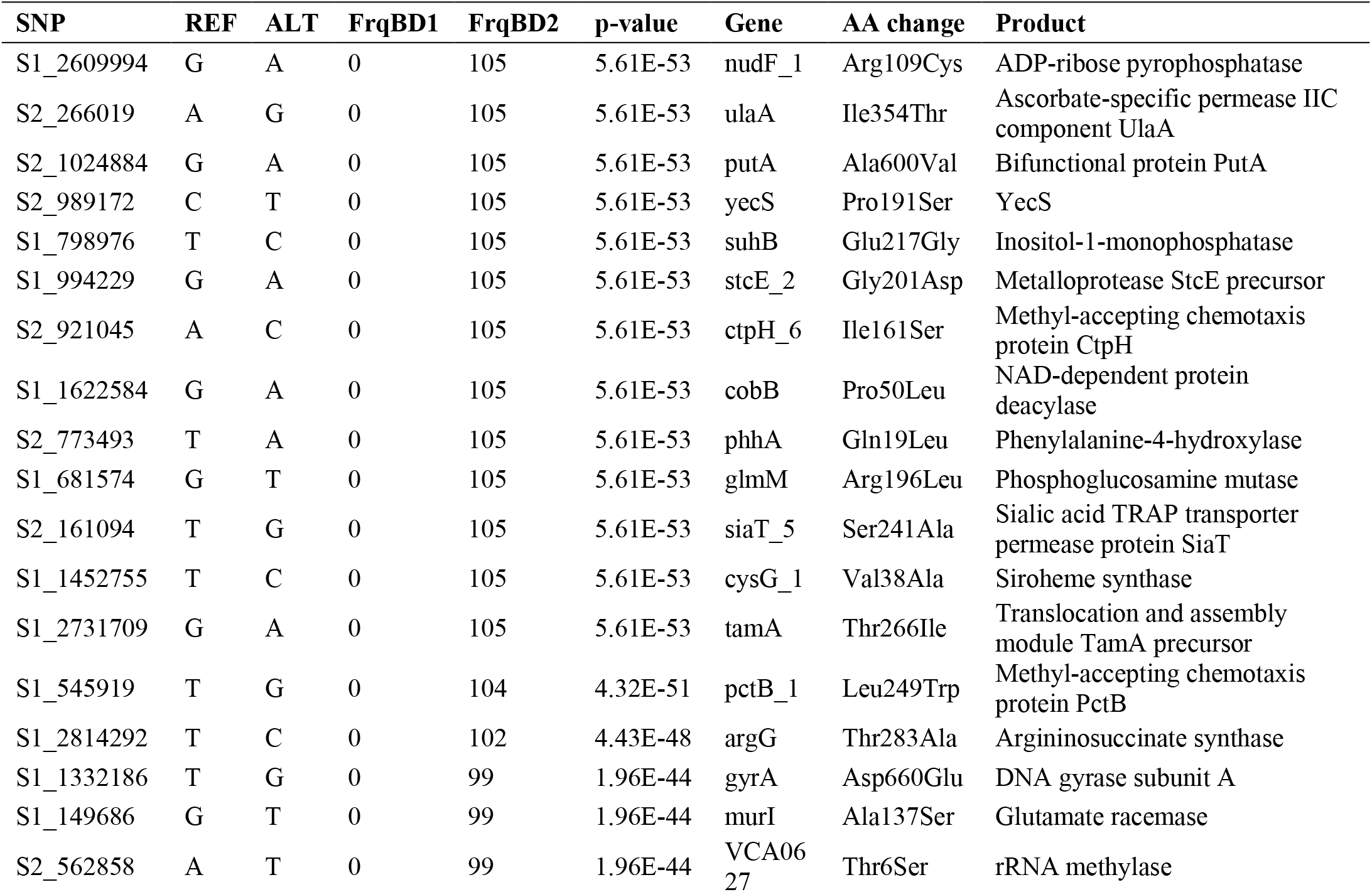
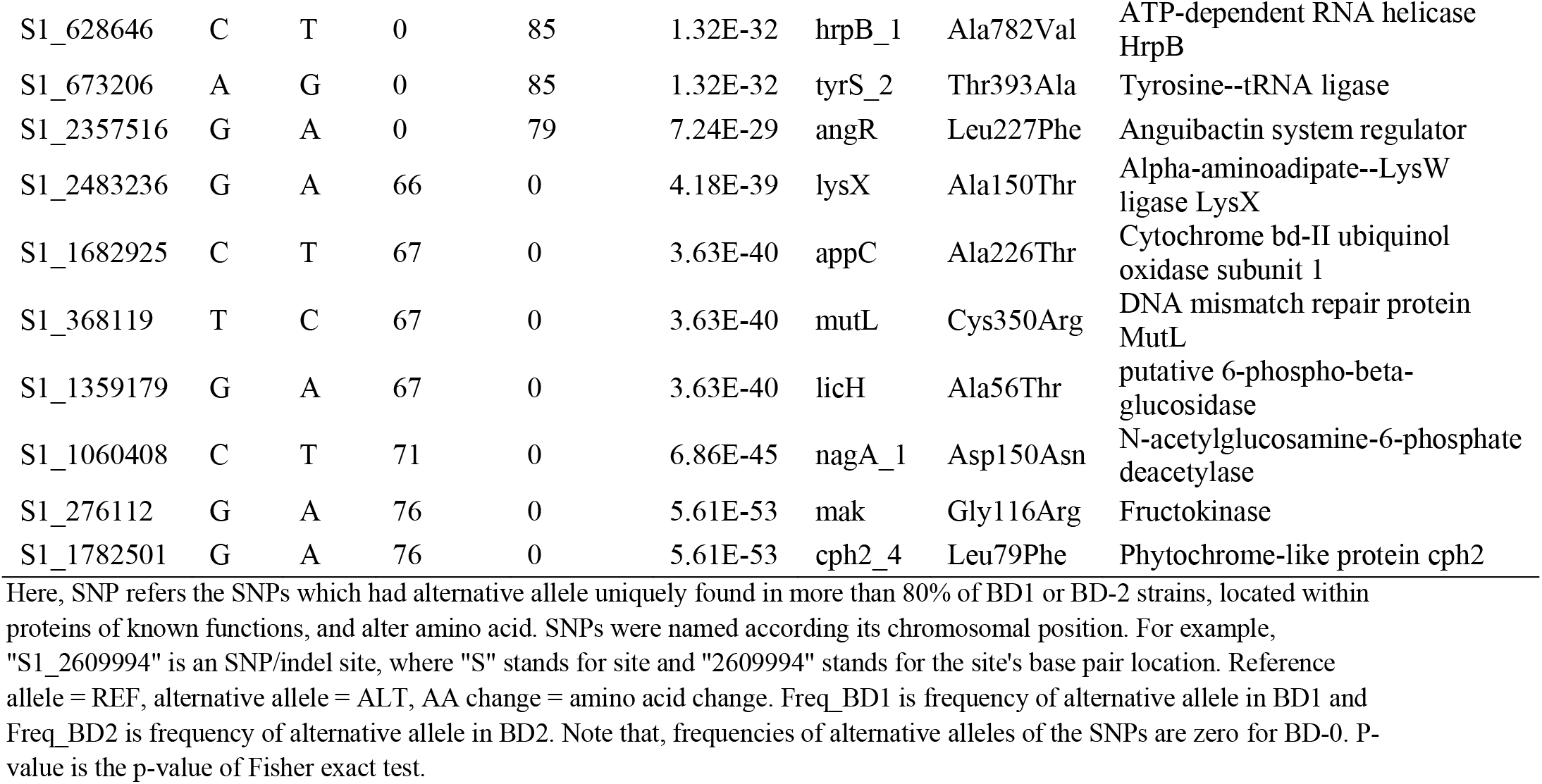
SNPs resulted unique mutant proteins in BD1 and BD2.

**FIG 3.**
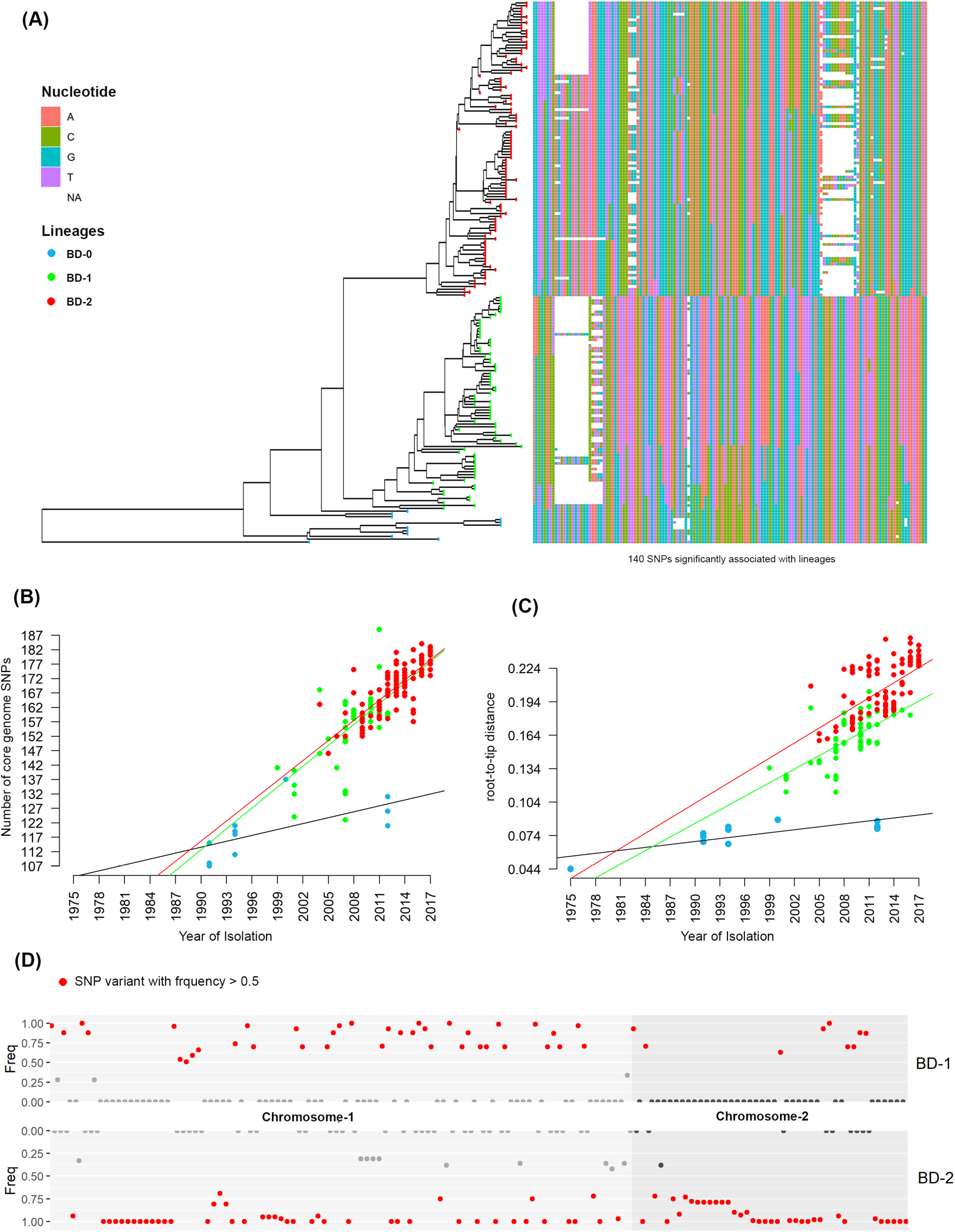
SNP analysis of genetic diversity. (A) Phylogenetic tree map of the strains and heat map for genotypes of 140 SNPs significantly associated with different lineages. Colors used delineate four different nucleotides where white represents the missing genotype. Heatmap shows clear differences in the lineages. (B) Number of core genome SNPs referencing the year of isolation. The figure shows steady accumulation of SNPs of different lineage strains over time. (C) Regression analysis of root-to-tip distance for strains of the lineages. This figure shows diversity of strains of different lineages. (D) Miami plot of alternative allele frequencies of SNPs for the dominant lineages BD-1 and BD-2. This figure shows the clear difference in SNP accumulation by the two dominant lineages BD-1 and BD-2.

### Relative gene abundance

Pangenome analysis was done using Roary to investigate differences in core and pan genes among the strains of BD-0, BD-1, and BD-2. Roary classified the identified functional genes into four categories: (i) core genes, present in 99-100% of the strains; (ii) softcore genes, present in 95-99% of the strains; (iii) shell genes, present in 15-95% of the strains; and (iv) cloud genes, present in less than 15% of the strains (22). Pangenome analysis revealed significant differences in overall gene composition among the clades (Fig 4A). According to the definition of core genes in pangenome analysis, the number of core genes largely varied among BD-0, BD-1, and BD-2 (see Table S4 in supplementary material). Similarly, the number of soft-core genes was also varied. BD-0 is a group of close relatives with a larger genetic distance relative to BD-1 and BD-2. All BD-0 strains and more than 95% of the BD-1 and BD-2 strains had 1102 common genes (see Table S5A in supplementary material) most having known function. About 10% of BD-2 strains had 44 unique genes of which six encoding crucial proteins of known function were found in more than 90% of the BD-2 strains. Those genes are: tetracycline repressor protein (**tetR**), tetracycline resistance protein (**tetA**), type-I restriction enzyme EcoKI M protein (**hsdM**), type-I restriction enzyme EcoR124II R protein (**hsdR**), Mrr restriction system protein (**mrr**), and 5-methylcytosine-specific restriction enzyme B (**mcrB**) (see Table S5B in supplementary material). In addition, methyl-accepting chemotaxis protein (**CtpH**) and group_10030 virulence proteins were exclusively found in 60% and 65% of BD-2 strains, respectively. By contrast, about 5-15% of the BD-1 strains carried 19 genes that were unique for them (see Table S5C in supplementary material). Three genes common to all BD-0 strains were not detected in BD-2 and were present only in 1-2 of the BD-1 strains.

**FIG 4.**
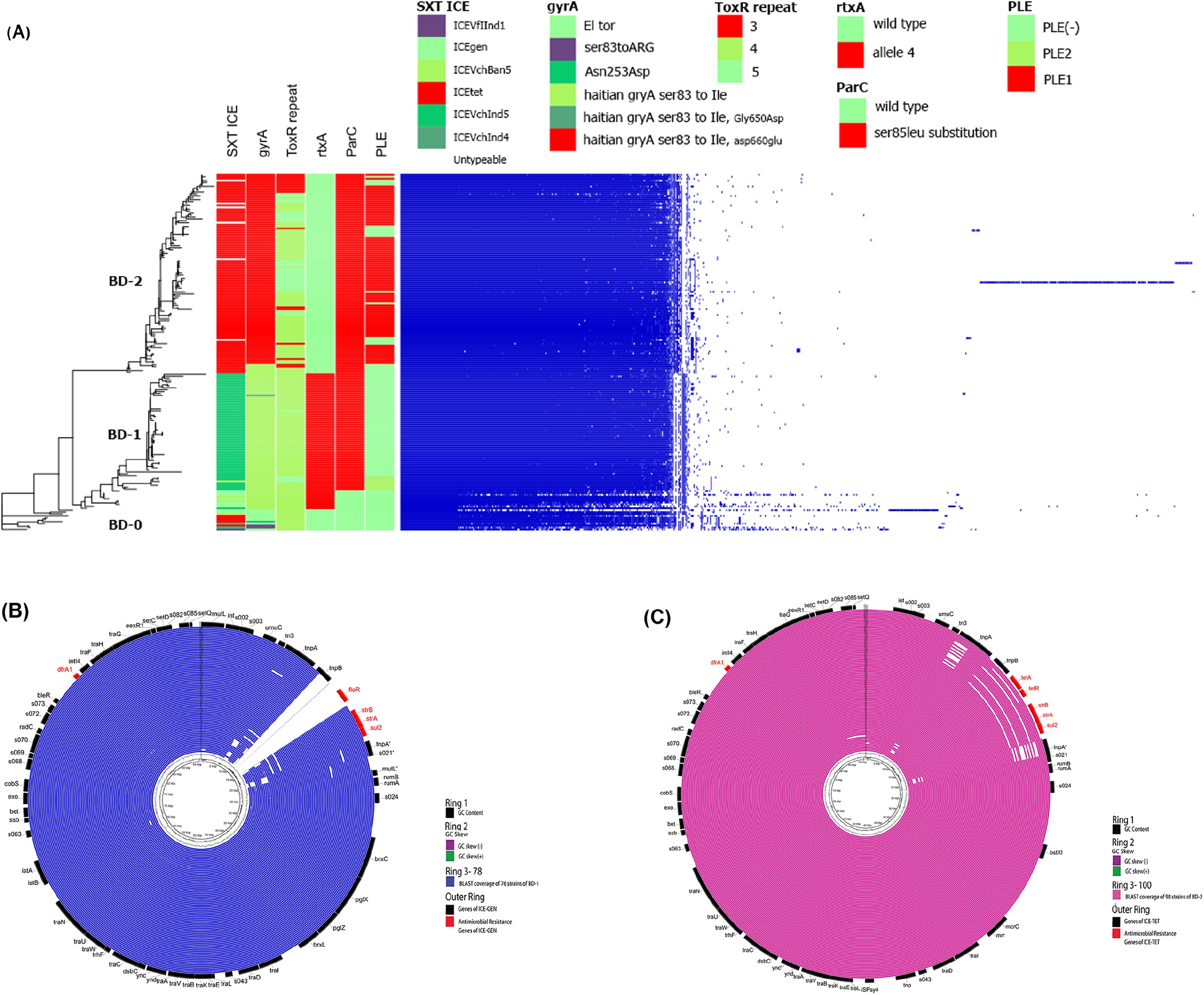
Pangenome analysis showing differences in abundance of gene clusters among the lineages. (A) Relative gene abundance of lineages identified by Roary. Features of the sequences are shown with bars and details for features listed in Table S1. (B) BLAST coverage of SXT regions of BD-1 isolates compared with ICE-GEN. Rings represent sequentially outwards following Table S1. Outermost ring shows the different genes of ICE-GEN. (C) BLAST coverage of SXT regions of BD-2 isolates compared with ICE-TET. The rings represent strains of BD-2 sequentially outwards following Table S1. The outermost ring shows different genes of ICE-TET.

Next, we conducted Pan-GWAS to identify clade-specific genes by considering gene presence and absence as the explanatory variable and defined lineage groups as the response variable. A total of 92 genes were significantly (p-value < 4.98×10^−6^) associated with BD-0 and BD-1 (see Table S6A in supplementary material). Of these, 62 genes were identified in 54-73% of BD-0 but not in BD-1 strains. Of 164 genes associated with BD-0 and BD-2, 46 were found in more than 73% of BD-2, but not in BD-0 strains (see Table S6B in supplementary material). In addition, 66 genes were found in more than 45% of BD-0, but not in BD-2 strains. Of 143 genes associated with BD-1 and BD-2 (see Table S6C in supplementary material), 29 were found in more than 76% of BD-1, but not in BD-2 strains. Again, 47 genes were found in 22-97% of the BD-2, but not in BD-1 strains. These results provide evidence that strains of BD-1 and BD-2 diverged and evolved as two lineages by accumulating genes, after originating from common ancestor BD-0.

### Pathogenicity islands and phage inducible chromosomal island like elements

*V. cholerae* strains included in this study were further examined by targeting the pandemic and pathogenicity islands namely VSP-1, VSP-II, VPI-1, and VPI-2, including the phage inducible chromosomal island like elements (PLE). Based on the extent of detected regions compared to *V. cholerae* N16961, five variants of VSP-II (variants 1-5 of the wild type) as reported in our recent study (16), and one variant of VPI-1 (variant 1 of the wild type) were observed (Fig 5). *V. cholerae* El Tor strains differed in type of VSP-II and VPI-1 variants. BD-0 had wild type of VSP-II, as in reference El Tor N16961 strain. Most BD-1 strains (except two) had variant-4 VSP-II, with partial deletion in VC_495 and complete deletion in VC_496 to VC_512, and BD-2 strains carried three VSP-II variants of which ca. 73% had variant-2 VSP-II with partial ORF VC_495 deletion, and complete VC_496 to VC_500 deletion, which appeared consistent with our prior study (16). BD-0 and BD-1 harbored wild type of VPI-1, whereas most of the BD-2 strains (102 of 105 strains) had variant VPI-1 with complete deletion of VC_819 to VC_820 ORFs; and partial deletion in VC_821. All BD-0 strains, and 66 of 76 BD-1 strains lacked PLE (see Tables S1 and S7 in supplementary material), while PLE2 was found in ten BD-1 strains isolated in 2007 possessing the *ctxB*1 genotype and one in 2005. Interestingly, most of the BD-2 strains (83 of 103) carried PLE1, but the rest lacked PLE. Thus, BD-2 lineage strains associated with recent Bangladesh endemic cholera are variant-3 VSP-II, variant VPI-1, and the majority possesses PLE1.

**FIG 5.**
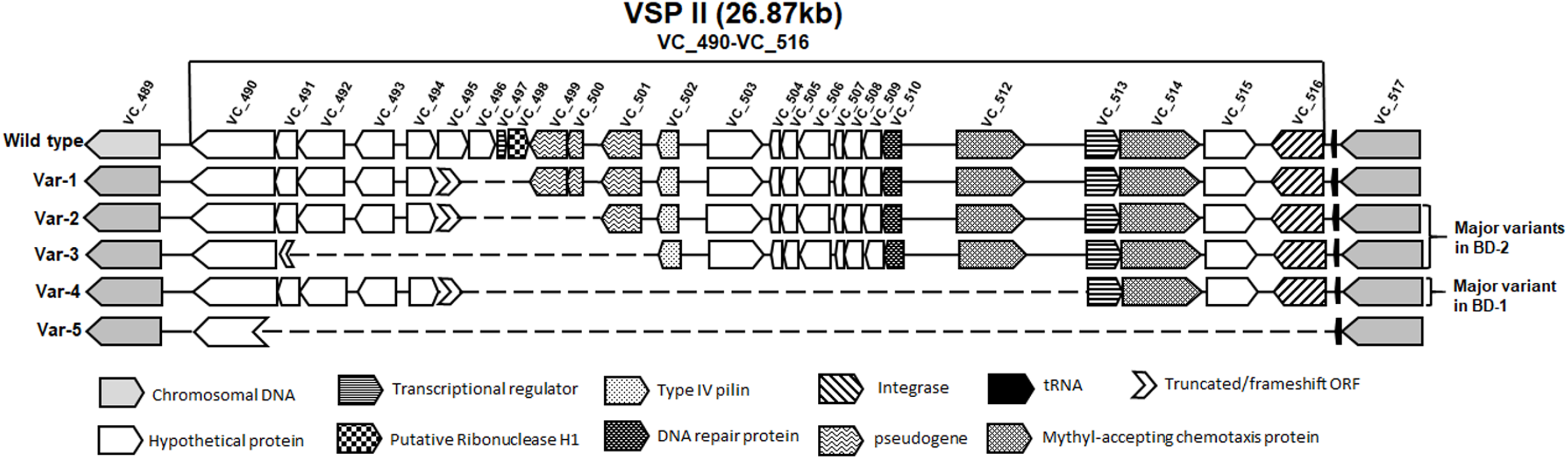
Schematic diagram of VSP-II. Schematic alignment view of VSP-II regions for the isolates. Direction of gene transcription is indicated by arrows and gene shadows represent functional annotation. Six types were identified with all BD-0 strains wild-type VSP-II. Two major types, var-2 and var-3, observed for most BD-2 strains and one major type var-4 for most BD-1 strains.

### Variations in SXT/R391 and important genes

Although differences in SXT/R391, *ctxB, gyrA, rtxA*, and *parC* across two lineages (BD-1, analogue of lineage-2; BD-2, analogue of lineage-1) were investigated in our recent study (16), these important genetic elements were rechecked to draw overall conclusions for all strains included in this investigation. Moreover, variation in ToxR binding repeats were checked across strains of different lineages. Integrative and conjugative elements (ICEs) were targeted from whole-genome sequences by aligning raw reads or contigs with five publicly available sequences of the ICE element (Accession ID: GQ463140.1, GQ463141.1, GQ463142.1, MK165649.1, and MK165650.1). Nucleotide blast was used to match extracted sequences with ICE element sequences and typed based on highest bit score. Four strains of BD-0 blast search yielded high bit scores when aligned with ICE^GEN^ (MK165650.1), *ICEVchInd5* (GQ463142.1), or ICE*Vch*Ban5 (GQ463140.1). Bit scores were highest for the other BD-0 strains when aligned with ICE^TET^ (Accession ID: MK165649.1), which has genomic characteristics similar to ICEVchVhn2255 (Accession ID: KT151660). For all BD-1 strain bit scores were high when aligned with ICE^GEN^, ICEVchInd5, or ICEVchBan5, and for BD-2 strains bit scores were highest when aligned with ICE^TET^, which is consistent with our previous results. All BD-1 and BD-2 strains contained mutant *gyrA* with an amino acid alteration Ser83Ile, whereas 99 (94.28 percent) of the 105 BD-2 strains exhibited Asp660Glu, which was not present in BD-1 or BD-0, also supporting our previous findings.

*V. cholerae* O1 El Tor strains in this study were CTX positive, and each carried a single copy of CTXΦ with a particular *ctxB* genotype. Three variants, *ctxB*1 (classical genotype), *ctxB*3 (typical El Tor genotype), and *ctxB*7 (Haitian variant), of the cholera toxin gene were detected and found associated with the clades (Fig. 1A). Similar to previous findings, all BD-2 strains had *ctxB*1 genotype, majority of BD-1 strains had *ctxB*7 genotype, and all but two BD-1 strains possessed *rtxA* that differed from El Tor reference N16961 by a single SNP at position 13602 of 1563748 bp (NCBI Accession ID: NC 002505.1), corresponding to *rtxA* allele 4 (23). However, in this study it was observed that early BD-1 strains had the *ctxB*1 genotype, and over time gained the *ctxB*7 genotype.

A prior study showed that, Kolkata strains had four heptad repeats (TTTTGAT), whereas Haitian strains had five heptad repeats (24). All BD-0 strains had four heptad repeats (Table S1), while most BD-1 strains (93.4%; n=71) had four repeats, and only 5.3% (n=4) strains had five repeats. As a result, the majority of BD-1 strains with *ctxB*7 genotypes differed from Haitian strains in ToxR binding repeats. BD-2 strains had more diversity in ToxR binding repeats with 59.0% (n=62) carrying heptad repeats, 24.8% (n=26) five repeats, and 16.2% (n=17) three repeats.

## Discussion

*Vibrio cholerae* biotype El Tor, the causative agent of the 7^th^ cholera pandemic has increased transmissibility and is more virulent than classical biotype (14, 15). The 7th pandemic strains of cholera circulating in Asia comprises two El Tor clades, one dominant in Bangladesh and the other in India (16). Genomic analyses that included additional strains and publicly available genome sequences of wave-2 and wave-3 strains (6, 12) provide a detailed view of longitudinally and temporally representative *V. cholerae* clades associated with endemic cholera in Bangladesh over a period of 27 years (1991 – 2017). The results provide new insights potentially interpretable as origin and progression, based on differences in SNPs, indels, and gene acquisition, including antibiotic resistance cassettes in BD-1 and BD-2, the latter having gained ascendency and dominance as the agent of Bangladesh endemic cholera.

Results of whole genome sequencing (16), combined with additional genome sequence data for *V. cholerae* El Tor isolates of Bangladesh endemic cholera, allowed identification of two lineages, designated BD-1 and BD-2. The two clades appear to have originated from a common ancestor of paraphyletic group BD-0, as early as 1981 (95% HPD: 1976-1986). According to A. Mutreja et al. (12), seven strains of BD-0 isolated between 1991 and 2000 represent wave-2 strains, and only one strain isolated in 1994, wave-3 with a most recent common ancestor (MRCA) for BD-1 and BD-2. The BD-1 and BD-2 clades may belong to wave-3. Although BD-0 consisted of predominantly of wave-2 strains, three sequenced strains isolated in 2012 shared a wave-2-like genetic background (6), suggesting wave-2 strains may have already been present. Almost all wave-3 strains from a previous study (12) grouped with strains belonging to BD-1. Consistent with results of a previous study (16), significant differences were noted between BD-1 and BD-2, which varied in temporal predominance as the causal agent of Bangladesh endemic cholera. Most (n=62; 82 percent) BD-1 strains had been isolated between 2007 and 2012, with predominance during that time. Between 2005 and 2017, 105 strains belonging to BD-2 were reported, with 97 obtained between 2009 and 2017, implying BD-2 association with recent Bangladesh endemic cholera until 2017. Phylodynamic analysis using BEAST (19) revealed strains of BD-1 had been isolated in Bangladesh roughly ten years before BD-2 strains (see, Fig. S1 in supplementary material), and previously identified as Asian lineage −2 and Asian lineage-1, respectively (16).

BD-1 and BD-2 strains appear to have advanced by accumulating different SNPs and indels. Fisher exact test (21) identified 140 SNPs and 31 indel differences between BD-1 and BD-2, resulting in gene alleles unique to them (Fig 3). The majority of the SNPs and indels were components of protein coding genes, suggesting a possible crucial role in their adaption in Bangladesh. Regression analysis of the number of SNPs and year of isolation suggested that both clades consistently accumulated SNPs over time, implying evolution in response to environmental selective pressure.

Pangenome analysis using Roary (22) provided evidence of gene acquisition by strains of the clades. A recent study of *V. cholerae* O1 strains isolated in Pakistan found evidence of gene acquisition, where the number of core and accessory genes varied among different lineages (25). According to results of the analysis reported here, the number of core and accessory genes varied significantly among strains of BD-0, BD-1, and BD-2 in Bangladesh (Fig. 4A). The Pan-GWAS approach helped identify genes unique for each clade which could be considered contributing to virulence and/or niche adaptation (26).

Phage inducible chromosomal island like elements (PLE) protect *V. cholerae* populations from ICP1 infection by acting as an abortive infection system (27). In this study, the observed predominance in BD-2 of PLE1, not found in BD-0 and BD-1, could have provided a selective advantage for the lineage over BD-1, establishing dominance as an etiological agent of endemic cholera in Bangladesh in recent years.

Two BD-0 strains carried CTX phage with *ctxB*3, while other strains carried CTX phage with typical *ctxB*1. Strains at the base of BD-1 had CTX with *ctxB*1 isolated before 2007 and comprised multiple clusters. Moreover, CTX phage of all BD-2 strains contained classical *ctxB*1. A mutation in *rtxA* creating a premature stop codon disabled toxin function in emerging *V. cholerae* El Tor strains bearing *ctxB*1 (24). As in the classical strains, altered El Tor pandemic strains eliminated *rtxA* after acquiring classical *ctxB*. In this study, BD-0 and BD-2 strains contained the wild-type *rtxA* allele 1 (Fig. 3A) described by Dolores and Satchell (23). None contained deletions in *rstB* gene when reads were compared to *V. cholerae* N16961 reference genome, indicating *rstB* of Bangladesh *V. cholerae* O1 El Tor isolates does not resemble that of the Haitian outbreak isolates that have been analyzed.

ToxR is a global transcriptional regulator of virulence gene expression and this repeated sequence is required for ToxR binding and activation of the *ctxAB* promoter. The ToxR-binding site is located immediately upstream of *ctxAB* and the affinity of ToxR binding is influenced by the repeat sequences (28). The presence of an increased number of ToxR binding repeats located between *zot* and *ctxA* has been hypothesized to correlate with a severe form of cholera (28). In this study, variation was detected in the number of ToxR binding repeats (TTTTGAT) among sequences of the *V. cholerae* El Tor isolates. All BD-0 strains had four heptad repeats observed in 93.4% of BD-1 and 59% of BD-2 strains. For BD-2 strains, however, greater variation was observed in ToxR binding repeats as ca. 24.8% (n=26) of BD-2 strains contained five heptad repeats, whereas 16.2% (n=17) had three heptad repeats, suggesting robustness of the clade.

Targets of quinolones are type II topoisomerases of DNA gyrase, a heterotetramer composed of two A and two B subunits, encoded by *gyrA* and *gyrB* genes respectively (29). It was observed that all BD-1 and BD-2 strains had a common mutation Ser83 to Ile in *gyrA*, while 94.29% (99/105) BD-2 had an additional mutation Asp660 to Glu. Furthermore, 87% (66/76) of BD-1 strains exhibited a mutation Ser85 to Leu *parC*, whereas all BD-2 strains (105/105) had this mutation. In Haitian *V. cholerae* strains, *gyrA* and *parC* genes had two point mutations: Ser83 to Ile in *gyrA* and Ser85 to Leu in *parC*. Both are linked to quinolone resistance in *V. cholerae* strains associated with recent cholera outbreaks in India, Nigeria, and Cameroon (30).

SXT/R391 family ICEs are transferable elements associated with antimicrobial resistance in *V. cholerae* (31). The SXT-ICE regions of the isolates included in this study, were compared with five sequences of the elements to the type SXT/R391 family ICEs belonging to strains associated with cholera (*V. cholerae* O1 and O139) (9, 32). Four BD-0 strains exhibited ICE elements similar to ICE^GEN^, ICEVchInd5 or ICEVchBan5, whereas the rest had ICE elements similar to ICE^TET^. Interestingly, ICE elements of BD-1 strains included ICE^GEN^, ICEVchInd5 or ICEVchBan5-like ICE elements, whereas BD-2 strains differed completely from the others, with only ICE^TET^-like ICE elements.

The results of the study reported here included BD-1 and BD-2 isolated during the Bangladesh endemic cholera of 2004 onwards and that, while existing together, with each subsequent year they exhibited different dominance. BD-2 diverged, while retaining the ability to produce multifunctional-autoprocessing repeats-in-toxin (MARTX) and acquiring SXT element ICE^TET^ containing tetracycline resistance genes. This observation hints at a selective advantage of BD-2 strains over BD-1 strains for robustness. It is evident from results of the analyses that BD-1 and BD-2 differ significantly, owing to gene composition and SNPs and may have evolved independently due to selection pressures. The use of antibiotics, including tetracycline, can exert selection pressure in evolution (16, 33), while strains stopping to produce MARTX along with other variations in the genome might provide a selective advantage. According to suggestions from studies of the dynamics of *V. cholerae*, immunocompetence of the host against *V. cholerae* strains may contribute to the dynamics of *V. cholerae*, hence produce an effect from interaction with humans in selection and cannot be ruled out (34).

Cholera globally is influenced by thriving populations of *V. cholerae* occurring naturally in the Ganges Delta of Bay of Bengal (GDBB) (1, 2, 5, 14). Overall results presented here suggest means of emergence and progression of the two clades in evolution from a progenitor *V. cholerae* El Tor initiating the seventh pandemic in Asia (5) and reflecting short-term evolution of *V. cholerae* El Tor associated with Bangladesh endemic cholera in the GDBB (14, 31). BD-2 is concluded to have emerged relatively recently and evolves by acquiring SNPs over time. Also, BD-2 strains showed diversity in indels, possessing SXT/R391 family ICE-elements, PLE1, *tetR*, and several other important genetic elements, and predominantly associated with recent Bangladesh endemic cholera. As is apparent from our results, BD-1 appears to be an analogue of a previously reported lineage 2 from Asia, the major causative agent of cholera in India, Yemen, and Haiti (16). In contrast, BD-2 strains of the present study appear to be an analogue of Asian lineage 1, which successfully outcompeted BD-1 (Asian lineage 2) and established predominance as an etiological agent of cholera in an historical hotspot of the disease, Bangladesh. It can be concluded that this is a reflection of robustness of BD-2 as an epidemic clone emerging locally with potential to transmit globally, and underscoring the need to track the two successful *V. cholerae* El Tor clades.

## Materials and Methods

### Bacterial isolates

A total of 119 *V. cholerae* O1 strains from the icddr,b collection of strains isolated in Bangladesh between 2004 and 2017 (see Table S1 in supplementary material) were sequenced. Paired-end Illumina short reads for the isolated strains were generated (150 bp, 150 bp) using MiSeq or Hiseq 2500 sequencer as described in our recent study (16). Publicly available paired-end raw reads of 17 strains isolated in Bangladesh between 1991 and 2007 (see study flow chart Fig. S3 in supplementary material) and 56 strains from our recent study (16) were included in the analysis.

### Genome assembly and gene annotation

An ultra-fast FASTQ preprocessor implemented in FASTP (35), was used to inspect raw paired-end reads and filter bad ligation or adapter parts. De novo genome assembly implemented in VelvetOptimizer (36) was used to build contigs by optimizing the parameter N50, a metric for assessing contiguity of an assembly. The bacterial genome annotation tool, Prokka (37), was used for whole-genome gene annotation. ResFinder (38) was used to find the antimicrobial resistant gene profiles for all of the strains.

### SNP identification and phylogenetic analysis

Bowtie2 (39) was used to align high-quality reads with reference genome sequence of *V. cholerae* N16961 El Tor (NCBI Accession ID: NC_002505.1 and NC_002506.1) for variant calling. Samtools (40) and Bcftools (41) were used to call genome variants. A maximum-likelihood phylogeny was inferred on an alignment of concatenated SNPs evenly distributed across non-repetitive, non-recombinant core genome using IQ-TREE v1.6.1 (42). Trees were visualized in FigTree v1.4.3 (http://tree.bio.ed.ac.uk/software/figtree/) or Interactive Tree of Life online tool (43).

### Bayesian phylogenetic inference

The Bayesian Evolutionary Analysis Sampling Trees (BEAST) v.2.4.4 software package (19) was used for temporal analysis to estimate divergence date of *V. cholerae* O1 isolates in Bangladesh. The date of isolation of each strain was used as tip data. A random clock model was implemented using Markov Chain Monte Carlo (MCMC) chains run for 100 million generations with 10% burn-in and sampled every 1000 generations. A GTR nucleotide substitution model was used. Tree data were summarized using TreeAnotator, a tool of BEAST software package, to generate the maximum clade credibility tree.

### Pangenome analysis

A pan-genome was constructed using Roary (22) from annotated assemblies of the sample set with percentage protein identity of 95%. The protein sequences were first extracted and iteratively pre-clustered with cd-hit (version 4.6) down to 98% identity. An all against all blast (version 2.2.31) was performed on the remaining non-clustered sequences and a single representative sequence from each cd-hit cluster was selected. The data were used by MCL (44) (version 11–294) to cluster the sequences. The preclusters and MCL clusters were merged and paralogs split by inspecting the conserved gene neighborhood around each sequence (5 genes on either side). Each sequence for each cluster was independently aligned using PRANK (45) (version 0.140603) and combined to form a multi-FASTA alignment of the core genes. Sequences of SXT elements were compared with ICE^GEN^ and ICE^TET^ using BRIG 0.95 with 70% BLAST identity (46).

## Acknowledgements

This work was supported in part by icddr,b, National Institutes of Infectious Diseases (NIID), Tokyo, and the Research Program on Emerging and Re-emerging Infectious Diseases (JP21fk0108139) from the Japan Agency for Medical Research and Development (AMED). Authors acknowledge icddr,b hospital and laboratory staffs for their support. icddr,b gratefully acknowledges the following donors for providing unrestricted support: Governments of the People’s Republic of Bangladesh, Global Affairs Canada (GAC), Swedish International Development Cooperation Agency (Sida), and the Foreign Commonwealth & Development Office (FCDO), UK. All the authors read and approved the final manuscript. None declared conflicts of interest.

## Supplementary Information

**FIG S1 Bayesian phylogenetic analysis of *V. cholerae* O1.** Node ages obtained from BEAST analysis. Tree visualized using FigTree v1.4.4. Colors of clades reference the lineage.

**FIG S2 Manhattan plots of *p*-values for association studies of SNPs and BD-1 and BD-2 lineages.** Blue represent suggested significant and red indicates high significance. Association analysis reveals 140 SNP difference between BD-1 and BD-2 lineages.

**FIG S3 Study flow chart.** Data curation and analyses steps are given in the flow chart.

**Table S1. Genetic characteristics of strains included in the study.** Lineage refers to genetically homogeneous groups of strains. Legends are strain ID, year of isolation, SXT/ICE elements, acquired antibiotic resistance profile, gyrA allele, number of ToxR binding repeats, ctxB allele, and PLE are tabulated.

**Table S2. Number of strains belonging to the different lineages.** Here, N_BD-0 = number of strains belonging to BD-0; N_BD-1 = number of strains belonging to BD-1; and N_BD2 = number of strains belonging to BD-2.

**Table S3. Fisher exact test identifying significantly associated 140 SNPs and 31 indels of the two dominant lineages, BD-1 and BD-2.** SNP/indel indicates significant SNP/indels identified by Fisher exact test. SNPs and indels named according to chromosomal position. For example, “S1 1905668” is an SNP/indel site, where “S” stands for site and “1905668” stands for location for site base pair. Reference allele is Ref, where as the alternative alleles are Alt1 and Alt2. P-value is Fisher exact test value. Variant type indicates SNP and indel type.

**Table S4. Roary pangenome analysis showing gene compositions differences by lineage.** Gene cluster refers to group of genes clustered based on existence in the strains of different lineages. Code for number of genes in lineage BD-0, BD-1, and BD-2 is N_BD-0, N_BD-1, and N_BD-2, respectively.

**Table S5. Common and unique genes of the different lineages.** (A) Genes detected in more than 95% of BD-0, BD-1, and BD-2 strains. (B) List of unique genes in BD-2. (C) List of unique genes in BD-1.

**Table S6**. **PanGWAS identified lineage associated genes.** (A) List of genes associated with BD-0 and BD-1. (B) List of genes associated with BD-0 and BD-2. (C) List of genes associated with BD-1 and BD-2.

**Table S7. Number of strains with phage inducible chromosomal island like elements (PLE)**. Absence of PLE = PLE(-), and PLE1 and PLE2 are two different types of PLE.

